# A new role for IFRD1 in regulation of ER stress in bladder epithelial homeostasis

**DOI:** 10.1101/2024.01.09.574887

**Authors:** Bisiayo E. Fashemi, Amala K. Rougeau, Arnold M. Salazar, Steven J. Bark, Rayvanth Chappidi, Jeffrey W. Brown, Charles J. Cho, Jason C. Mills, Indira U. Mysorekar

## Abstract

A healthy bladder requires the homeostatic maintenance of and rapid regeneration of urothelium upon stress/injury/infection. Several factors have been identified to play important roles in urothelial development, injury and disease response, however, little is known about urothelial regulation at homeostasis. Here, we identify a new role for IFRD1, a stress-induced gene that has recently been demonstrated to play a critical role in adult tissue proliferation and regeneration, in maintenance of urothelial function/ homeostasis in a mouse model. We show that the mouse bladder expresses IFRD1 at homeostasis and its loss alters the global transcriptome of the bladder with significant accumulation of cellular organelles including multivesicular bodies with undigested cargo, lysosomes and mitochondria. We demonstrate that IFRD1 interacts with several mRNA-translation-regulating factors in human urothelial cells and that the urothelium of *Ifrd1^−/−^* mice reveal decreased global translation and enhanced endoplasmic reticulum (ER) stress response. *Ifrd1^−/−^* bladders have activation of the unfolded protein response (UPR) pathway, specifically the PERK arm, with a concomitant increase in oxidative stress and spontaneous exfoliation of urothelial cells. Further, we show that such increase in cell shedding is associated with a compensatory proliferation of the basal cells but impaired regeneration of superficial cells. Finally, we show that upon loss of IFRD1, mice display aberrant voiding behavior. Thus, we propose that IFRD1 is at the center of many crucial cellular pathways that work together to maintain urothelial homeostasis, highlighting its importance as a target for diagnosis and/or therapy in bladder conditions.

## INTRODUCTION

The urinary bladder performs the primary function of storing and releasing urine and is lined with a pseudo-stratified transitional layer, i.e., urothelium, which forms a highly specialized impermeable barrier and is composed of three main cell types (superficial, intermediate, and basal). The superficial cell layer harbors uroplakin proteins that localize to the superficial apical plasma membrane and help form the impermeability barrier and protect from toxic factors in urine ^1^. Under homeostatic conditions, the urothelium is quiescent and has the slowest turnover of any mammalian epithelia ^2–4^. However upon injury, there is a rapid activation of regenerative response leading to complete restoration within 72 hours post injury ^5–10^. A number of factors have been identified to play important roles in urothelial development, injury and disease response ^11–14^.

The bladder undergoes distension-contraction cycles as it fills and voids throughout each day and across the whole lifespan. The vast changes in apical membrane surface area required by voiding cycles depend on superficial cells’ intracellular discoidal fusiform vesicles lined with four uroplakin proteins (Ia, Ib, II, and III) that are recruited to the apical surface during distension ^15–19^. Upon contraction, these uroplakin proteins are then recycled via endocytosis and multivesicular bodies (MVB) trafficking to decrease the size of the urothelial membrane; damaged uroplakins can also be routed from MVBs to lysosomes for degradation ^20–23^. Thus, the urothelium is exposed to urinary wastes and must withstand shear forces from continual expansion and contraction. The molecular mechanisms underpinning how the urothelium maintains homeostasis under this state of constant stress is not well understood.

The interferon-related developmental regulator 1 (IFRD1, a.k.a PC4 and TIS7), is a protein whose exact molecular function remains unknown ^24–26^, but it has recently been shown to play a significant role in paligenosis, a cellular dedifferentiation process key for mediating injury response in the pancreas, stomach, liver, and kidney. Paligenosis (for *pali*, return, + *gene*, generative, + *osis,* a process) is an evolutionarily conserved cellular program that facilitates recruitment of differentiated cells into the cell cycle after injury to regenerate damaged cells ^27,28^. ^29^. Both transcription and translation of IFRD1 are stimulated by tissue and cell injury, including endoplasmic reticulum (ER) stress; indeed, *IFRD1* mRNA is one of a handful of transcripts translated even when cellular stress has caused global decrease in translation via phosphorylation of the translation-regulating protein eIF2α ^30,31^. *Ifrd1^−/−^*mice are fertile and viable, and young adults have not been noted to have phenotypes at homeostasis in multiple organs.

In this study, we show that, surprisingly, IFRD1 is expressed robustly in the urinary bladder at homeostasis (i.e,. in the absence of injury), and, remarkably, loss of function caused significant molecular, cellular, and organellar alterations in *Ifrd1^−/−^*urothelium. Cytological and immunological assays revealed significant increase in cargo-filled multivesicular bodies (MVBs), lysosomes, and mitochondria, and abnormal distribution of uroplakin in *Ifrd1^−/−^* urothelium. Transcriptomic analysis revealed loss of IFRD1 led to substantial alterations, including changes in expression of genes encoding pathways regulating function of mitochondria, ER, and the cell cycle. We also show that loss of IFRD1 significantly impacted global translation levels in the bladder with increased ER stress and activation of the unfolded protein response (UPR). Moreover, loss of IFRD1 enhanced the production of reactive oxygen species (ROS), increased cell death markers, and increased epithelial cell shedding into the urine. Finally, we note a decreased urothelial regenerative capacity and aberrant voiding behavior in *Ifrd1^−/−^* mice. Overall, our data demonstrate the importance of IFRD1 in maintaining bladder homeostasis.

## RESULTS

### IFRD1 is expressed in bladder at homeostasis and loss of IFRD1 causes alterations in urothelial cellular architecture and molecular pathways

Young (8-12 weeks old) wild-type (WT) mouse bladders showed robust expression of IFRD1 protein **(Figure 1A, lane 1)**, whereas, as expected, *Ifrd1^−/−^*mice did not have detectable IFRD1 **(Figure 1A, lane 2)**, supporting our use of this mouse model to dissect the role of IFRD1 in bladder homeostasis. To assess how the loss of IFRD1 affects the global environment of the bladder at homeostasis, we performed total RNA Sequencing (RNAseq) on bladder tissue from young WT and age-matched *Ifrd1^−/−^* mice. In comparison with WT bladders, *Ifrd1^−/−^* bladders displayed at least 1.5-fold (FDR-adjusted P<0.05) decrease in expression of 53 genes and increase in expression of 161 genes **(Figure 1B)**. Gene set enrichment analysis of this altered bladder transcriptome using the MSigDB Collections identified statistically significant downregulation of 8 pathways and upregulation of 20 in *Ifrd1^−/−^*mice **(Figure 1C).** Notably, the downregulated pathways included protein secretion, reactive oxygen species, oxidative phosphorylation, fatty acid metabolism pathways **(Figure 1C)**, all of which are critical for urothelial function. To determine whether these transcriptional alterations were correlated with histological changes associated with the bladder tissue, we performed H&E staining of the unperturbed WT and *Ifrd1^−/−^* urothelium **(Figure 1D)**. *Ifrd1^−/−^* bladders were notable for harboring luminal cellular debris, and urothelial cells showed a marked and consistent increase in intracellular vesicles **(Figure 1D, bottom panel)**. Transmission electron microscopy (TEM) revealed accumulation of multivesicular bodies (MVBs) **(red arrow)**, as recognized by their characteristic, vesicle-within-vesicle morphology in *Ifrd1^−/−^* superficial cells **(Figure 1D)**. While the MVBs in WT superficial cells had lumens with the characteristic electron-lucent (“empty”) or amorphous material, the MVBs of *Ifrd1^−/−^* superficial cells were packed with undigested cargo **(Figure 1E, right panel, white arrows and inset)**. While we did not observe a significant difference in the amount of total MVBs, we found that there was a statistically significant difference in the census of cargo-filled MVBs within the *Ifrd1^−/−^* superficial cells **(Figure 1F)**. Additionally, *Ifrd1^−/−^* superficial urothelial cells harbored more numerous mitochondria **(black arrow) (Figure 1G)** and lysosomes than **(Figure 1H)** versus WT cells.

**Figure 1.**
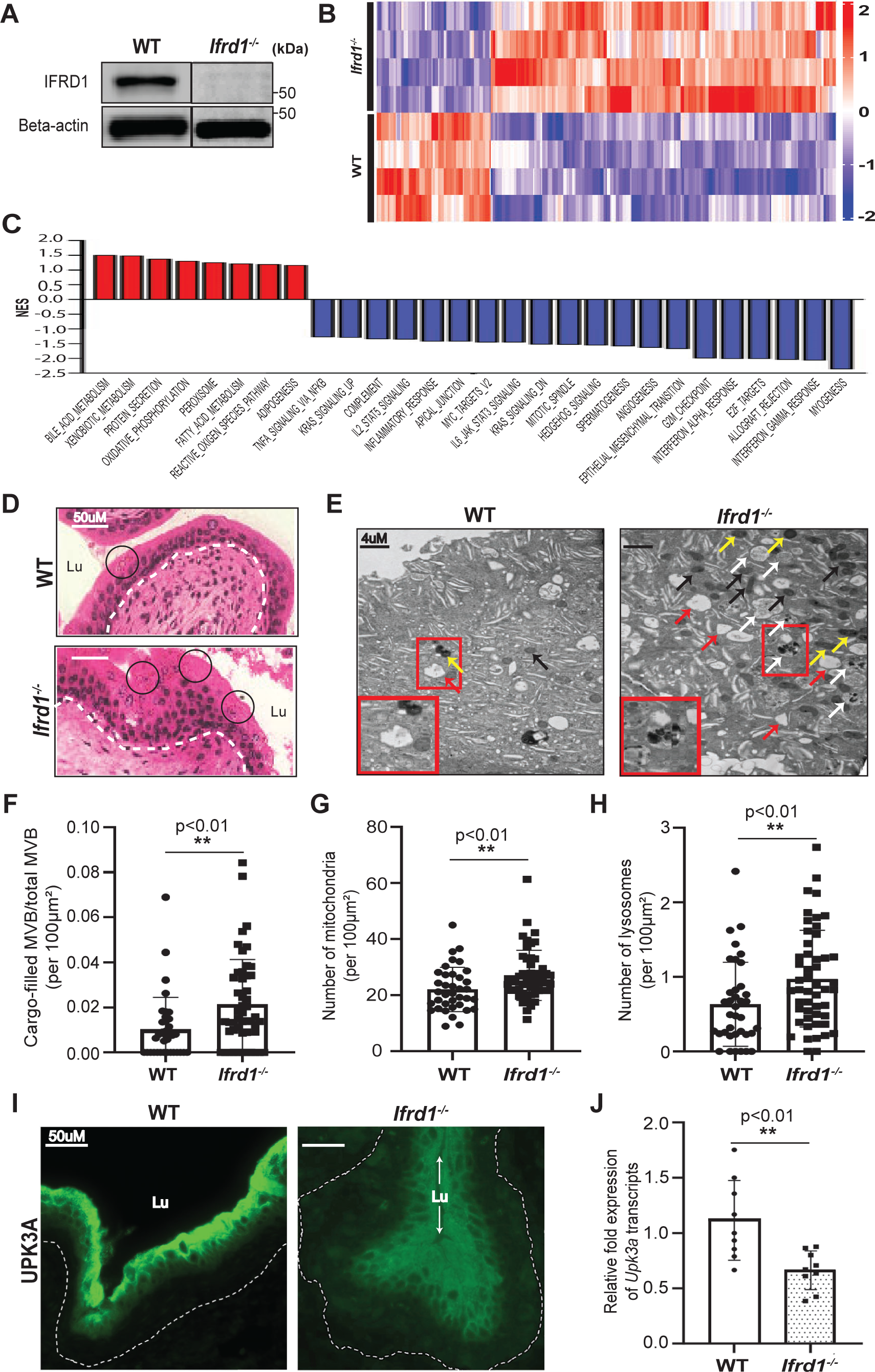
IFRD1 is expressed in bladder at homeostasis and loss of IFRD1 shows alterations in urothelial cellular architecture and molecular pathways. (A) Western blotting (WB) of wild-type (WT) and *Ifrd1* null (*Ifrd1^-/-^*) bladders at homeostasis. Beta-actin serves as a housekeeping control. (B) Heatmap of the global tissue transcriptome of WT and *Ifrd1^-/-^* bladders with fold change >1.5 and Benjamini-Hochberg FDR-adjusted p <0.05 (n = 4 bladders/group). (C) GSEA Hallmark pathway analysis showing significantly downregulated (red bars) and upregulated (blue bars) pathways in *Ifrd1^-/-^* bladders with respect to WT, based on fold change expression of related genes. (D) H&E staining show vesicles in WT (highlighted in black circles) and accumulated vesicles in *Ifrd1^-/-^* bladders in the superficial cells. (E) Transmission electron microscopy (TEM) of superficial cells show MVB (red arrows) in WT and *Ifrd1^-/-^*. MVBs in *Ifrd1^-/-^* show accumulated undi-gested material (white arrows). *Ifrd1^-/-^* superficial cells also display more lysosomes (yellow arrows) and mitochondria (black arrows) compared with WT. Magnified insets show the empty (WT) and cargo-filled (*Ifrd1^-/-^*) MVBs. (F) Analysis of cargo-filled MVB to total MVB ratio per 100μm*^2^* in WT and *Ifrd1^-/-^* superficial cells (Mann-Whitney test, unpaired non-para-metric test). (G) The number of mitochondria per 100μm*^2^* in WT and *Ifrd1^-/-^* superficial cells (Mann-Whitney test, unpaired non-parametric test). (H) The number of lysosomes per 100μm*^2^* in WT and *Ifrd1^-/-^* superficial cells (Mann-Whitney test, unpaired non-parametric test). (I) Immunostaining of UPK3A (green) in WT and *Ifrd1^-/-^* superficial cell layer (n= 6 blad-ders/group). (J) Real-time quantitative PCR (RT-qPCR) analysis of *Upk3a* transcripts in WT and *Ifrd1^-/-^* bladders (Mann-Whitney test, unpaired non-parametric test).

Given that the *Ifrd1^−/−^* superficial cells displayed an overabundance of MVBs with undigested cargo, we wanted to test whether loss of IFRD1 affected uroplakin trafficking. As expected, uroplakin III (UPIII) staining in the WT was expressed along the apical surface of the urothelium **(Figure 1I, left panel)**, In contrast uroplakins were diffusely distributed within superficial cells in *Ifrd1^−/−^* bladders **(Figure 1I, right panel)**, indicating aberrant uroplakin trafficking. Accordingly, transcripts encoding UPIII were significantly decreased in *Ifrd1^−/−^* bladders **(Figure 1J)**. Together, these findings demonstrate that IFRD1 is expressed at bladder homeostasis, and its loss is associated with significant cellular and alterations resulting in gross defects in the urothelial ultrastructure even in an unperturbed state.

### IFRD1 interacts with mRNA-translation-regulating proteins, and its absence is associated with global translational changes and enhanced ER stress

To identify the molecular partners that IFRD1 may be affecting or interacting with in the urothelium, we performed proteomic analysis of the urothelial cell line 5637 to capture the IFRD1 interactome at homeostasis. We performed co-immunoprecipitation followed by tandem mass spectrometry (MS/MS) using an antibody that specifically recognizes IFRD1 and a non-specific, IgG control, **(Figure 2A)**. We identified a total of 2271 proteins that interacted with IFRD1 or the IgG control at a probability rate of over 95%. Of those 2271 protein hits, 444 were detected in precipitates using IgG alone, 943 using IFRD1 alone, and 884 were found in both IFRD1 and IgG precipitates **(Figure 2B)**. For our analysis, we focused on the hits that exclusively interacted with IFRD1.The 943 IFRD1-only hits were ranked using their Total Spectrum Count. We then selected the interactors with 2 or more unique peptides and FDR <0.05. This resulted in a total of 584 unique proteins **(Figure 2C)**. These proteins were then subjected to PANTHER Overrepresentation test for GO molecular function. Excitingly, a large number of these proteins were found to be enriched in regulating mRNA translation (e.g., eukaryotic translation initiation factors [eIFs], mRNA binding proteins, preribosome binding factors) **(Figure 2C).** These molecular findings together with our cellular observations of accumulated MVBs with undigested proteinaceous cargo, suggested that IFRD1 might affect protein translation.

**Figure 2:**
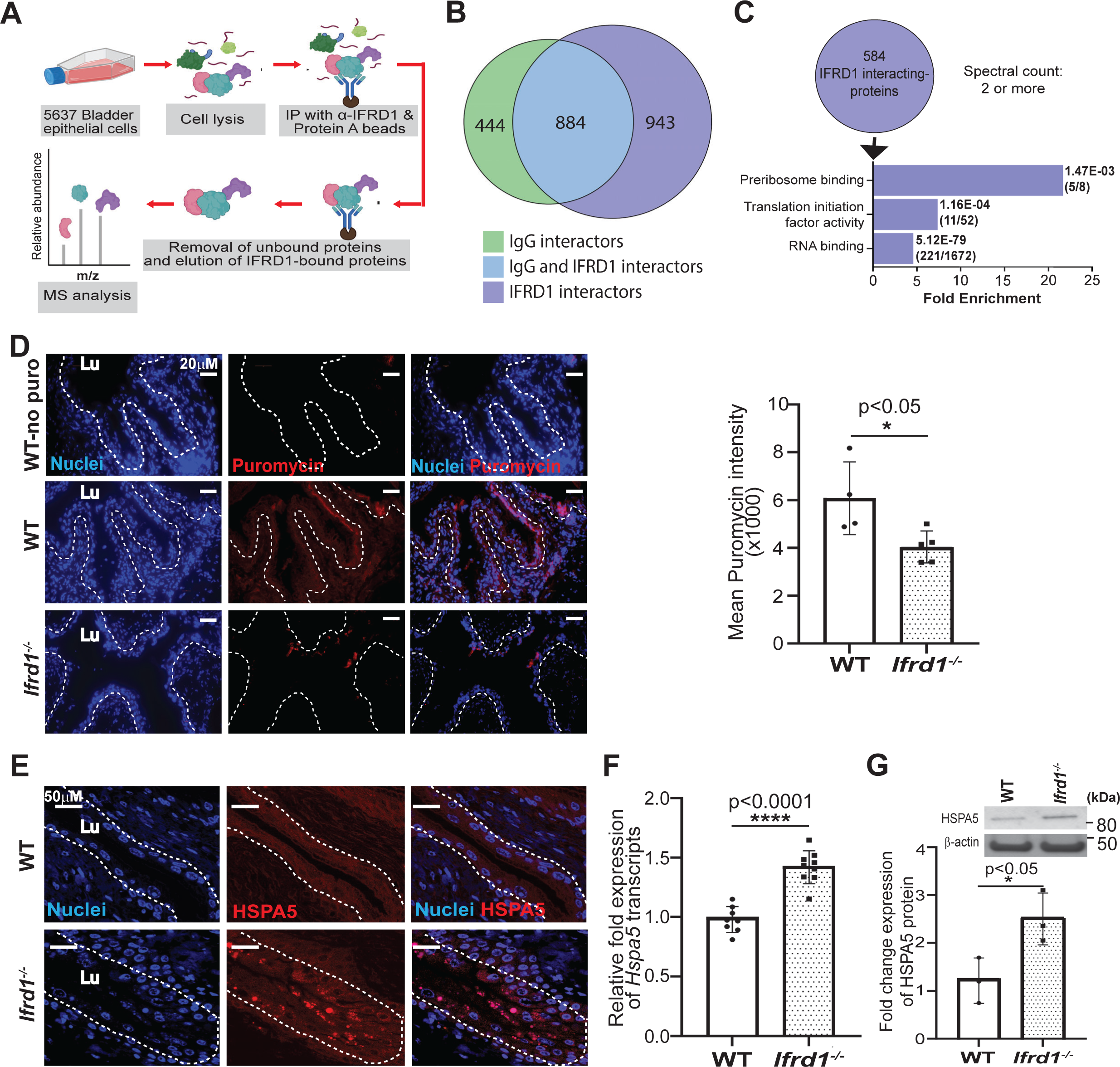
IFRD1 interacts with proteins involved in regulation of RNA translation and its absence is associated with global translational changes and enhanced ER stress. (A) Schematic diagram of co-immunoprecipitation of IFRD1 with tandem mass spectrometry analysis in 5637 bladder epithelial cell line. **(B)** Venn diagram shows the relationship of the protein interactors between IgG antibody control and IFRD1 pulldowns. **(C)** Fold enrichment plot of the GO Molecular Functions of a subset of IFRD1 interactors with 2 or more spectral counts. FDR values and fraction (in parenthesis) indicated are of those that matched with the reference list in PAN- THER database. **(D)** Immunostaining of bladders shows the expression of puromycin (red) in WT and *Ifrd1^-/-^* urothelia (WT-no puro represents the negative control where mice did not receive puromycin treatment prior to freezing). Nuclei are stained with DAPI (blue) (n= 3 mice/group). Quantitation of puromycin signal intensity (Data presented as mean ± SD, n= 4, each point is an average of 3 ROIs from different images, p values by two-tailed unpaired t test) **(E)** Immunostaining of HSPA5 (red) in WT and *Ifrd1^-/-^* urothelia. Nuclei are stained with DAPI (blue) (n= 3/group). **(F)** RT-qPCR analysis of *Hspa5* transcripts (n=8/group, p value by Mann-Whitney test) **(G)** WB and densitometric quantitation of HSPA5 protein levels. Beta-actin serves as a housekeeping control (Data presented as mean ± SD, n = 3/group, p value by one-tailed Mann-Whitney test).

To test whether the loss of IFRD1 would influence global protein translation levels in the bladder, we performed SunSET (surface sensing of translation) assay utilizing puromycin, an inhibitor of translation elongation, that can measure changes to protein synthesis *in vivo*, as it incorporates into elongating polypetide chain and induces translation termination ^32^. We injected WT and *Ifrd1^−/−^* mice with puromycin for 2 hours to allow the incorporation of puromycin into actively translating polypeptides. Immunofluorescence analysis showed that global translation in the *Ifrd1^−/−^* urothelium was decreased versus WT bladders **(Figure 2D, E)**.

Protein translation is a tightly regulated process in the maintenance of proteostasis and alterations are associated with an overabundance of abnormal, misfolded proteins within the endoplasmic reticulum (ER) leading to an ER stress response. To test whether such ER stress response is associated with the observed changes in translation levels in *Ifrd1^−/−^* bladders, we first investigated levels of the ER chaperone, BiP (HSPA5), which is upregulated under conditions of ER stress. We found that the *Ifrd1^−/−^* urothelium showed increased BiP in superficial cells **(Figure 2E)**. And BiP abundance in *Ifrd1^−/−^* bladders bladder was increased at both transcript **(Figure 2F)** and protein level **(Figure 2G)**. Thus, the results revealed that IFRD1 interacts with proteins involved in RNA translation, and *Ifrd1^−/−^* bladders display decreased global protein translation and enhanced ER stress response.

### Enhanced ER stress associated with loss of IFRD1 is linked with activation of the PERK arm of the UPR pathway

Next, we sought to determine how IFRD1 was effecting the ER stress response. ER stress can be induced via the unfolded protein response (UPR) activated by three ER-transmembrane, upstream stress sensors: ATF6 (activating transcription factor 6), IRE1α (inositol-requiring enzyme 1α), and PERK (protein kinase R-like endoplasmic reticulum kinase) ^33,34^. Under homeostatic conditions, BiP is bound to the ER-transmembrane stress sensors proteins, rendering these proteins inactive ^33,34^. However, unfolded, ER-associated proteins can bind BiP, dissociating BiP from the sensors. Each sensor, once no longer BiP-bound, can then activate its signature pattern of downstream gene expression changes **(schematized in Figure 3A)**.

**Figure 3.**
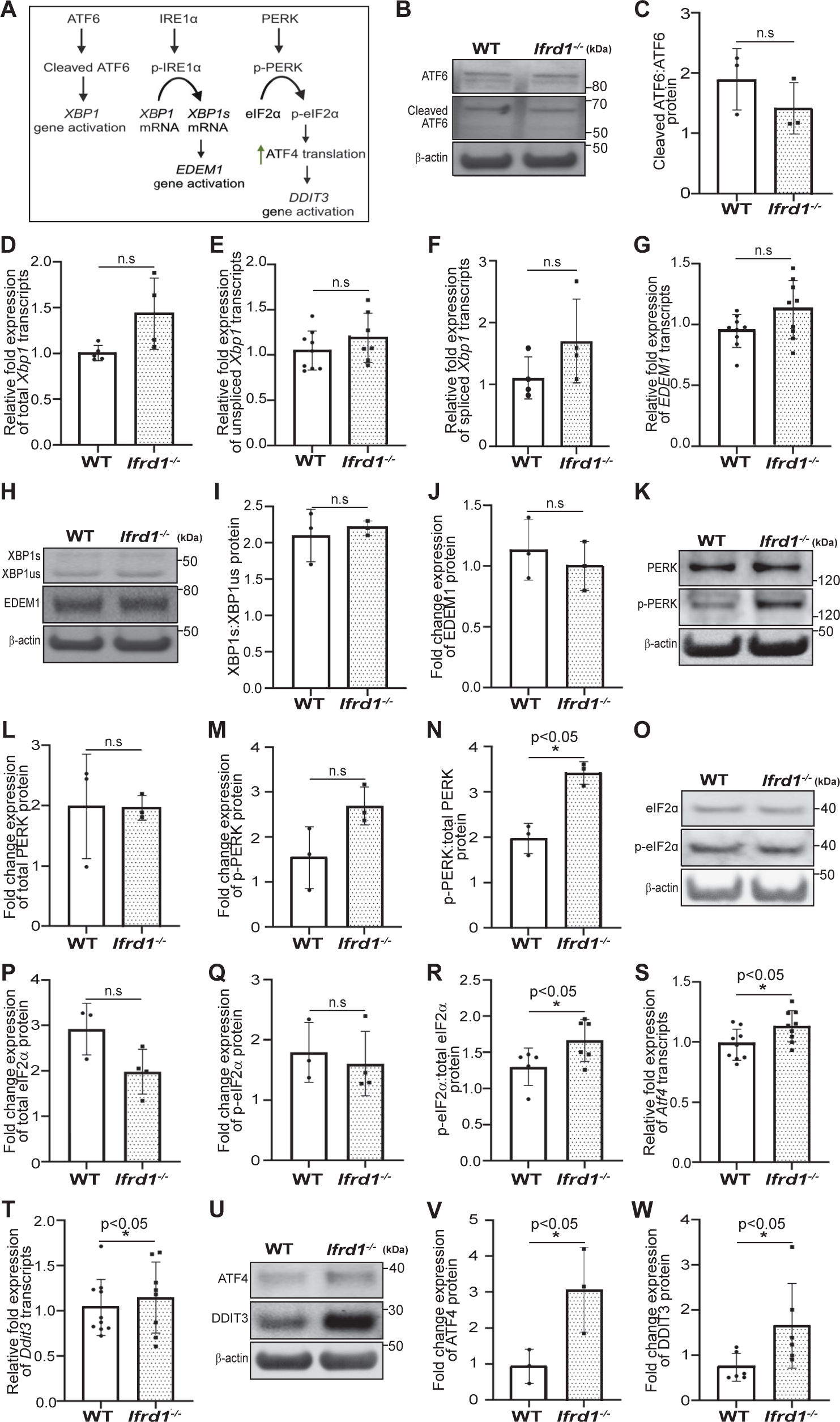
Enhanced ER stress associated with loss of IFRD1 is linked with activation of the PERK arm of the UPR pathway. (A) The three arms of unfolded protein response. **(B)** WB of full-length and cleaved ATF6 proteins in WT and *Ifrd1^-/-^* bladders. Beta-ac-tin was used as a loading control. **(C)** Densitometric quantitation of the ratio of cleaved ATF6 to total ATF6 protein levels (Data presented as mean ± SD, n = 3/group, p values by one-tailed Mann-Whitney test). **(D)** RT-qPCR analysis of total *Xbp1* transcript levels. (Data presented as mean ± SD, n = 5 WT, 4 *Ifrd1^-/-^* bladders, p values by one-tailed Mann-Whitney test) **(E)** RT-qPCR analysis of *Xbp1us* (unspliced) transcript levels (Data presented as mean ± SD, n = n = 8 WT, 7 *Ifrd1^-/-^* bladders, p values by one-tailed Mann-Whitney test). **(F)** RT-qPCR analyses of *Xbp1s* (spliced) transcript levels (Data presented as mean ± SD, n = 4/group, p values by one-tailed Mann-Whitney test). **(G)** RT-qPCR analyses of *Edem1* transcript levels (Data presented as mean ± SD, n = 9/group, p values by one-tailed Mann-Whitney test). **(H)** WB of XBP1s, XBP1us, and EDEM1 proteins in WT and *Ifrd1^-/-^* bladders. Beta actin was used as a loading control. **(I)** Densitometric quantitation of the ratio of XBP1s to XBP1us protein levels (Data presented as mean ± SD, n = 3/group, p values by one-tailed Mann-Whitney test). **(J)** Densitometric quantitation of EDEM1 protein levels (Data presented as mean ± SD, n = 3/group, p values by one-tailed Mann-Whitney test). **(K)** WB of PERK and p-PERK proteins in WT and *Ifrd1^-/-^* bladders. Beta-actin was used as a loading control. **(L)** Densitometric quantitation of total PERK (Data presented as mean ± SD, n = 3/group, p values by two-tailed Mann-Whitney test). **(M)** Densitometric quantitation of p-PERK (Data presented as mean ± SD, n = 3/group, p values by two-tailed Mann-Whitney test). **(N)** Densitometric quantitation of the ratio of p-PERK to PERK (Data presented as mean ± SD, n = 3/group, p values by one-tailed Mann-Whitney test). **(O)** WB of e F2a and p-e F2a prote ns n WT and *Ifrd1^-/-^* bladders. Beta-actin was used as a loading control. **(P)** Dens tometr c quant tat on of total e F2a (Data presented as mean ± SD, n = 3 WT, 4 *Ifrd1^-/-^* bladders, p values by two-tailed Mann-Whitney test). **(Q)** Dens tometr c quant tat on of p-e F2a (Data presented as mean ± SD, n = 3 WT, 4 *Ifrd1^-/-^* bladders, p values by two-tailed Mann-Whitney test). **(R)** Dens tometr c quant tat on of the rat o of p-e F2a to e F2a (Data presented as mean ± SD, n = 5 WT, 6 *Ifrd1^-/-^* bladders, p values by one-tailed Mann-Whitney test). **(S)** RT-qPCR analysis of *Atf4* transcript levels (Data presented as mean ± SD, n = 9/group, p values by two-tailed Mann-Whitney test). **(T)** RT-qPCR analy-sis of *Ddit3* transcript levels (Data presented as mean ± SD, n = 10 WT, 8 *Ifrd1^-/-^* bladders, p values by one-tailed Mann-Whitney test). **(U)** WB of ATF4 and DDIT3 proteins in WT and *Ifrd1^-/-^* bladders. Beta-actin was used as a loading control. **(V)** Densitometric quantitation of ATF4 (Data presented as mean ± SD, n = 3/group, p values by one-tailed Mann-Whitney test). **(W)** Densitometric quantitation of DDIT3 (Data presented as mean ± SD, n = 6/group, p values by one-tailed Mann-Whitney test).

We systematically examined impact of loss of IFRD1 on each of the arms of the UPR by analyzing the signature downstream targets. First, we examined ATF6. When ATF6 is released from BiP, it translocates to the Golgi where it is cleaved to release its N-terminal active form, ATF6-N (cleaved ATF6)^35^. We determined abundance of total (p90) and cleaved (p50) ATF6. We saw no significant ATF6 activation in *Ifrd1^−/−^* bladders **(Figure 3B-3C**, one-tailed t-test). RT-qPCR analysis of its downstream transcriptional target, *Xbp1* similarly showed no significant increase in expression **(Figure 3D)**.

Next, we determined impact of loss of function of IFRD1 on IRE1α which, on release of BiP, homodimerizes and autophosphorylates leading to induction of its RNase activity to splice the *Xbp1* unspliced mRNA (*Xbp1us*) to *Xbp1 spliced mRNA* (*Xbp1s*), an active form that encodes the XBP1 protein isoform that can translocate into the nucleus to induce expression of genes such as *Edem1* ^36^.. We analyzed the levels of *Xbp1us* and *Xbp1s* variants transcript and protein levels via RT-qPCR **(Figure 3E-3F)** and WB **(Figure 3H top panel, 3I)**, respectively and saw no significant increase in *Ifrd1^−/−^* bladders, consistent with IRE1α signaling not being IFRD1-dependent. Further, the XBP1s gene target, *Edem1*, also showed no significant increase in transcript **(Figure 3G)** or protein **(Figure 3H middle panel, 3J)** abundance in mouse bladders lacking IFRD1 in comparison with WT.

Finally, we determined whether IFRD1 affects the PERK arm, which similarly to IRE1α, homodimerizes and autophosphorylates upon BiP release. Activated PERK then phosphorylates a protein that regulates translation initiation, eIF2α. Phosphorylated eIF2α decreases activity of the eIF2 ternary complex (eIF2-GTP-tRNAi) that is required for the translation initiation ^37,38^. Consistent with the PERK arm being activated by loss of IFRD1, we observed a significant increase in the ratio of phosphorylated to total PERK in the *Ifrd1^−/−^* bladders compared to that of WT bladders **(Figure 3K-3M**, one-tailed t-test). If PERK were activated in the absence of IFRD1, we would also expect an increase in the phosphorylation of p-PERK target, eIF2α. Indeed, our WB analysis revealed a significant increase in the ratio of phosphorylated to total eIF2α protein levels **(Figure 3O-3R**, one-tailed t-test). p-PERK phosphorylation of eIF2α leads to global protein translation suppression while favoring translation of select transcripts such as *Atf4*, which induces expression of genes like *Chop* (*Ddit3*) ^33,34,39^. Accordingly, both *Atf4* and its target *Chop* were upregulated significantly at both transcript and protein level **(3S- V)** in the *Ifrd1^−/−^* bladders. Together, our results indicate that loss of IFRD1 in mouse bladder specifically affects the activation of PERK arm of the UPR which may a mediator of the urothelial stress response.

### *Ifrd1^−/−^* mice display increased urothelial cell death, dysregulated renewal and aberrant voiding behavior

We wondered whether there would be physiological relevance of the observed UPR activation of PERK-ATF4-CHOP pathway in *Ifrd1^−/−^* bladders. As unresolved stress can lead to persistent activation of CHOP which can in turn lead to apoptosis when stress is not resolved, we examined whether the *Ifrd1^−/−^*urothelial cells exhibit increased apoptosis. To that end, TUNEL assay, which fluorescently labels cells positive for double-stranded DNA breaks that are generated during continued cells stress and apoptosis showed near universal positivity in *Ifrd1^−/−^* mice with rare to no positive cells in WT bladder **(Figure 4A)**. Persistent ER stress/cell death is associated upstream and downstream with increase in ROS. To test whether this is true for *Ifrd1^−/−^* bladders, we stained the urothelium with the ROS indicator dye Dihydroethidium (DHE) which showed stronger staining as corroborated by the higher fluorescence intensity in *Ifrd1^−/−^* bladders **(Figure 4B)**. We next examined if the constitutively enhanced ROS and apoptosis levels altered the urothelial tissue homeostasis in *Ifrd1^−/−^*bladders. TEM images **(Figure 4C, TEM)** and cytological examination of urines (**Figure 4C, Pap stain**) from WT and *Ifrd1^−/−^* mice revealed a significant level of spontaneously shed epithelial cells in *Ifrd1^−/−^*urine samples. Quantification of the shed cells by Pap stain revealed about 4-fold increase in the *Ifrd1^−/−^* urine samples at baseline with many shed cells also showing strong ROS signal **(Figure 4C, D)**. Thus, not only did the *Ifrd1^−/−^* bladders shed a significantly higher number of cells, but they were positive for ROS.

**Figure 4.**
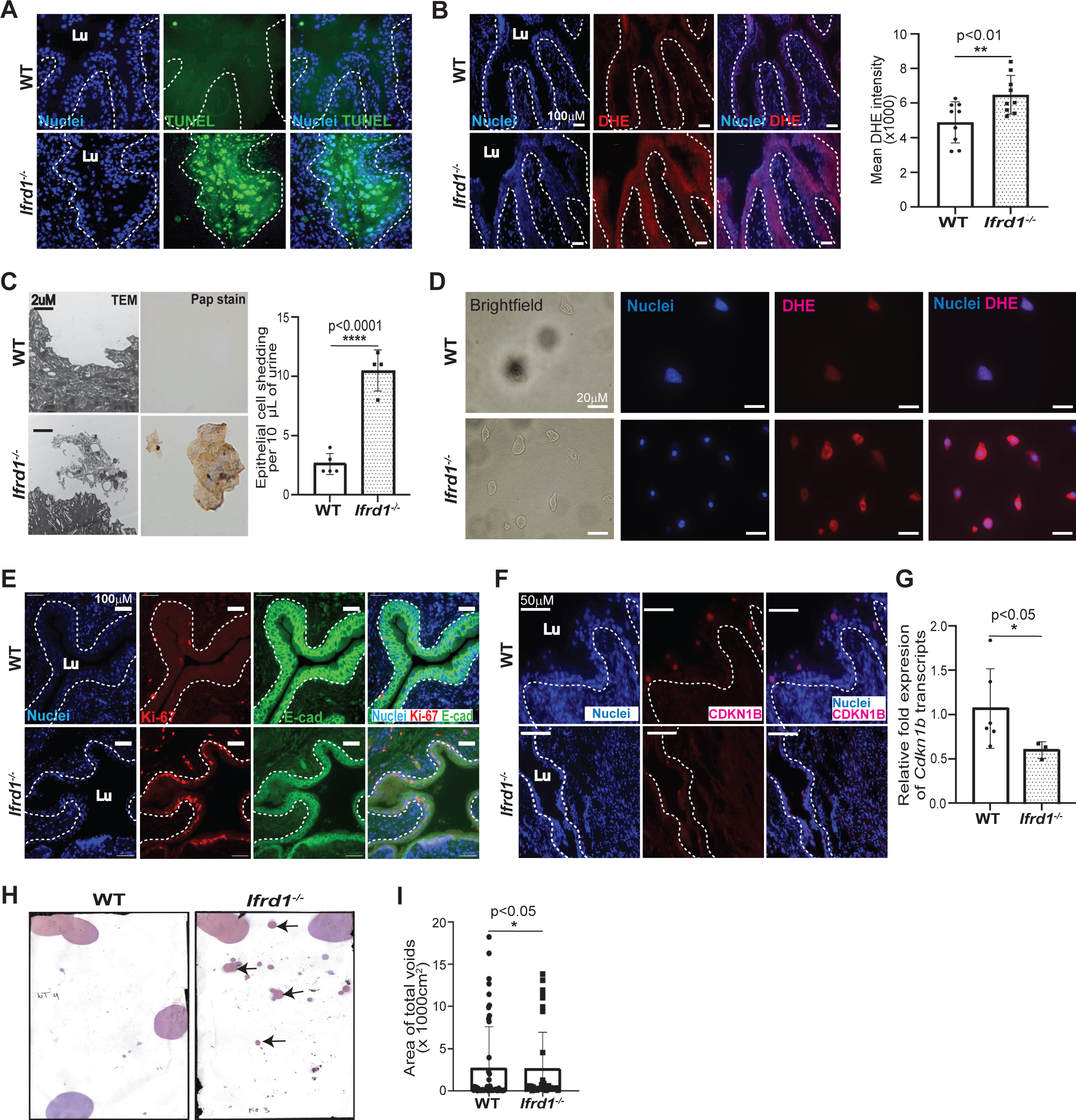
Ifrd1 null mice display increased urothelial cell death, dysregulated renewal and aberrant voiding behavior. (A) TUNEL staining of WT and *Ifrd1^-/-^* urothelial cells (n= 3/group). **(B)** DHE staining of WT and *Ifrd1^-/-^* urothelia. Quantitation of DHE staining intensity (Data presented as mean ± SD, n= 9, 3 ROIs from 3 different images per group, p values by two-tailed unpaired t test). **(C)** Representative TEM (left) and urine cytology (right) images show exfoliated bladder epithelial cells. Quantitation of exfoliated/shed epithelial cells (Data presented as mean ± SD, n= 5 WT, 6 *Ifrd1^-/-^* mice, p values by two-tailed unpaired t test). **(D)** Brightfield and immunostaining for DHE (red) in exfoliated epithelial cells of WT and *Ifrd1^-/-^* bladders. Nuclei are stained with DAPI (blue) (n= 3/group). **(E)** Immunostaining for Ki-67 (red), E-cadherin (green) in WT and *Ifrd1^-/-^* urothelial cells (n= 3/group). **(F)** Immunostaining of CDKN1B (red) in WT and *Ifrd1^-/-^* urothelia (n= 3/group). **(G)** RT-qPCR analysis of *Cdkn1* transcripts in WT and *Ifrd1^-/-^* bladders (Data presented as mean ± SD, n= 5 WT, 6 *Ifrd1^-/-^* bladders, p values by Mann-Whitney test. **(H)** Representative images and **(I)** quantitation of void spot assay of WT and *Ifrd1^-/-^* mice (n=8, p-values by Mann-Whitney test).

Under homeostatic conditions, the virtually quiescent urothelium contains few proliferatively active cells, which reside within the basal cell compartment (^4,8,9,40,41^. Urothelial shedding in response to unmitigated ROS could represent a stunted response to repair damaged/stressed cells which often leads to compensatory proliferation in the basal cell layer of the *Ifrd1^−/−^* urothelium. As expected, we found that WT bladders contained only scattered Ki67-positive cells **(Figure 4E, top)**. However, within the *Ifrd1^−/−^* urothelium, we found virtually every basal cell expressed Ki67 **(Figure 4E, bottom)**. Additionally, E-Cadherin staining, which typically labels cell-cell junctions in basal and intermediate urothelial cells, was reduced in the *Ifrd1^−/−^* urothelium, further indicating that urothelial integrity is constitutively compromised in these mice **(Figure 4E)**.

Normally, increased proliferation of basal cells fuels subsequent differentiation of these cells to regenerate superficial cells. Surprisingly, however, expression of p27KIP1 (CDKN1B), a marker for terminally differentiated superficial cells, was decreased in p27kip1+ superficial cell nuclei in the *Ifrd1^−/−^* urothelium **(Figure 4F)**, and *p27kip1* transcript abundance was also decreased overall in *Ifrd1^−/−^* bladders **(Figure 4G)**. These observations suggest that the loss of IFRD1 at homeostasis is not only linked to increased ER stress response and cell death but also with impaired regeneration of superficial cells, potentially compromising urothelial function in storing and voiding of urine. Therefore, we next sought to examine the voiding behavior of the WT and *Ifrd1^−/−^* mice. When placed on a filter paper, mice typically seek edges to urinate. However, *Ifrd1^−/−^*mice voided all along the filter paper and on the edges **(Figure 4H, black arrows)** with increased overall area covered by urine hinting that the *Ifrd1^−/−^* mice suffer incontinence **(Figure 4I)**.

## DISCUSSION

The urothelium is remarkable in its flexibility and impermeability. Injury such as that induced by infection, or mechanical/chemical stressors in the tissue results in rapid epithelial exfoliation and renewal and restoration of homeostasis. In the current study, we identify a new role for the transcription factor, IFRD1(which has largely been studied in tissues for its role in response to cell injury or in aging) in preventing spontaneous induction of the bladder injury response. We show that mouse bladder not only expresses IFRD1 at homeostasis but also that its loss is associated with alterations in urothelial morphology and function. We show that IFRD1 associates with proteins involved in protein translation, and *Ifrd1^−/−^*bladders/urothelium had reduced global translation with concomitant activation of the ER stress response. ER stress in the absence of IFRD1 was linked to activation of the PERK arm of unfolded protein response (UPR), and the *Ifrd1^−/−^* bladders exhibited increased oxidative stress, cell death and reduced terminal differentiation and aberrant voiding behavior. Thus, our new work suggests that even homeostatic activity of the bladder urothelium could constitute a potential ‘stress’ situation that requires management of UPR/ER stress by proteins such as IFRD1 that are normally required only in stressful situations in other tissues.

IFRD1 is an immediate-early gene induced by a variety of signals/stresses implicated in regulation of growth and differentiation of neurons, myocytes, enterocytes ^42–45^, and elevated levels of IFRD1 are observed in multiple acute injury models such as muscle trauma, small intestine resection, ischemia-reperfusion, stroke ^44,46–48^ highlighting a key role for IFRD1 in mediating cellular and tissue regeneration after injury. Our work on paligenosis reiterated the early activation of IFRD1 following injury, and its absence led to decreased regeneration and increased cell death in diverse organs such as stomach, liver, kidney, and pancreas in mice and human. Additionally, injury-induced expression of IFRD1 is found to be conserved in other organisms such as Axolotls and *Schizosaccharomyces pombe* ^27,49^. Given such critical roles of IFRD1 in regulating injury response, it is surprising that IFRD1 plays a seemingly steady-state role in the bladder. We speculate that perhaps the continual voiding cycles of the urinary bladder may require extensive trafficking and recycling of uroplakins that produces a consistent stressed state, requiring the urothelium to constitutively express stress-relieving proteins.

Injury and infections are known to engage inflammatory pathways in response to stress, and a growing number of studies point to the role of UPR activation to alleviate such stress and restore homeostasis ^50,51^. However, when UPR fails to do this, ER functions such as protein-folding, processing and trafficking are impaired for an extended period, which can induce apoptosis. Our results demonstrating a diffuse expression pattern and reduced expression of the uroplakin, UPK3A, along with increased TUNEL staining, support the conclusion that UPR is constitutively activated, causing both uroplakin trafficking defects in the bladders of IFRD1 null mice as well as constant pro-apoptotic signaling. We note that TUNEL simply labels DNA strand breaks, which usually correlate with – but are not entirely specific – for apoptosis. In terminally differentiated cells like those of the superficial urothelium, it is possible that strand breaks accumulate as a result of ROS (which are also elevated in the absence of IFRD1) and that repair is slow in these non-dividing cells, even if they are not all targeted for eventual death. We also noticed not only decreased p27kip1 in the bladder in the absence of IFRD1 but also a diffuse, intracellular distribution of this normally nuclear protein, suggesting altered proteostasis of this protein as well as decreased differentiation. The increased ROS and other aberrant cellular features correlated with epithelial cell shedding in the urine of *Ifrd1^−/−^* mice. Moreover, we notice that the frequent shed cells in these mice had smaller nuclei than those of WT suggesting that some of these shed *Ifrd1^−/−^* cells were less differentiated intermediate cells, which are not normally seen in urine. Urine usually contains only rare shed superficial cells, unless the bladder encounters stresses such as infection ^8,52,53^. Thus, *Ifrd1^−/−^* mice may experience cell death and shedding of cells even prior to terminal differentiation, correlating with the reduction in p27kip1 staining and expression.

The increased shedding and other cellular abnormalities in *Ifrd1^−/−^* mice are reminiscent of recent work ^54^ from our group demonstrating that aged bladders (>15 months of age) show enhanced DNA damage, increased ROS, and higher levels of epithelial shedding. In other words, loss of IFRD1 might trigger an aging-like phenotype. It may be that these effects noted in aging are partly mediated via the functions of IFRD1 and its association with translation, ER stress and UPR activation. In ongoing studies, we will determine if IFRD1 expression decreases with aging and if that might be a molecular basis for the aging phenotype. Additionally, IFRD1 has been shown to exert some of its effects via modulation of transcriptional activity via its interaction with histone deacetylases (HDACs) and its altered expression is associated with loss of polarity in epithelial cells via its interaction with the HDACs ^55–57^. Indeed, our MS/MS analysis revealed IFRD1 interaction with HDACs (HDAC4 and SIRT6, data not shown) in the urothelium. As altered polarity in the epithelial cells is linked with lower E-Cadherin expression and cancer including bladder cancer ^58^, we could speculate that alterations to the E-Cadherin levels we note in the urothelium upon loss of IFRD1 may have a more profound impact as the mice get older.

Our work here hints that the seemingly quiescent urothelium is under a constant state of stress owing to the massive structural transitions it undergoes daily during normal voiding behaviors and that IFRD1 plays a key role in modulating the ER stress response and maintaining the urothelial homeostasis. Loss of IFRD1 appears to alter voiding dynamics. While our limited observations do not explain why or how, it is nevertheless striking that stress urinary incontinence (SUI) and pelvic organ prolapse (POP), conditions associated with voiding dysfunction have been linked to ER stress and POP has even been correlated with altered IFRD1 expression in women^59,60^. Specifically, postmenopausal women with SUI have shown an activated PERK arm of the UPR with no changes to IRE1α or ATF6 arms, resembling the phenotype observed in IFRD1 null mice from our study^60^. Additionally, this study showed increased apoptosis mediated by PERK-ATF4-CHOP correlates with the SUI in postmenopausal women. Similarly, *IFRD1* mRNA levels were altered in POP tissues compared to controls in modulating the pathogenesis of POP^59^. Given our findings that IFRD1 affects ER stress with a physiologically relevant phenotype that is reproduced in women with bladder dysfunctions, studies are warranted to determine the involvement of aberrant UPR in POP-SUI pathology and to delineate the sequence of events caused by the absence of IFRD1 leading up to the SUI pathology highlighting its importance in diagnosis and/or therapy ^61^.

Overall, our findings suggest that IFRD1 is a new player with a critical role in the maintenance of urothelial structure and bladder functions at homeostasis and its loss is associated with alterations to proteostasis and impaired superficial cell renewal with pathophysiologic results on bladder function. Understanding the mechanisms of action of IFRD1 may shed light on urothelial homeostatic activities that are altered in the aged or injured/infected/diseased bladders.

## AUTHOR CONTRIBUTIONS

BEF, AK, AMS, CC, JCM and IUM conceived the experimental plan; BEF and AK performed most experiments assisted by AMS, RC, SB. JWB and CC provided expertise. BEF, AK, AMS, JCM, and IUM wrote the manuscript, and all authors approved the final draft.

## ACKNOWLEDGMENTS

This work was supported in part by NIH grants, R01DK100644, R01AG052494, P20DK119840, and R56AG064634 (to IUM), T32-AI007172 (to BEF); by NIH grants, K08DK132496, R21 AI156236, P30 DK052574, Department of Defense grant through the PRCRP program under Award No. W81XWH-20-1- 0630, and by the Doris Duke Charitable Foundation Fund to Retain Clinical Scientists (to JWB); Department of Defense, through the Peer Reviewed Cancer Research Program (PRCRP) program under award W81XWH2210327 (to CJC); by NIH R01DK105129, R01CA239645, and P30 DK056338 (to JCM). Thanks also to Ilja Vietor and Lukas A Huber, Division of Cell Biology, Biocenter, Medical University of Innsbruck, Innsbruck, Austria, for the original generation of *Ifrd1^−/−^* mice. We also thank the Tissue Analysis and Molecular Imaging Core of the Texas Medical Center Digestive Disease Center for histological processing (supported by P30 DK056338) and to Dr. Wandy Beatty of the Washing University Microscopy Score.

## DECLARATION OF INTERESTS

IUM serves on the scientific advisory board of Luca Biologics. No conflicts of interest exist.

## STAR METHODS RESOURCE AVAILABILITY

### Lead contact

Further information and requests for resources and reagents should be directed to and will be fulfilled by the lead contact, Indira U. Mysorekar.

## Materials availability

This study did not generate new unique reagents.

## Data and code availability

- This study does not report original code.
- Data are available on the Gene Expression Omnibus under the following identifier: GSE149571.

## EXPERIMENTAL MODEL AND STUDY PARTICIPANTS

### Mice

C57B6/J mice (12-16 weeks: WT) were obtained from the mouse facility at Washington University School of Medicine and Baylor College of Medicine. Standard rodent chow and water were available ad libitum throughout the experiment. *Ifrd1^−/−^* mice were a kind gift from Dr. Lukas Huber (Medical University Innsbruck) and Dr. Deborah Rubin (Washington University School of Medicine). Mice were housed in groups of four to five in a temperature-(22 ± 1 °C) and humidity-controlled vivarium with lights maintained on a 12:12 light/dark cycle. All animal experimental procedures were approved by the Institutional Animal Care and Use Committee at Washington University School of Medicine (Animal Welfare Assurance #A- 3381-01) and Baylor College of Medicine (Animal protocol number AN-8629). All mice were humanely euthanized at the end of each experiment.

## METHOD DETAILS

### Cell lines

Human urinary urothelial carcinoma cell line (5637, HTB9) (ATCC) was cultured in RPMI 1640 media with 10% fetal bovine serum in a humidified atmosphere at 37 °C with 5% CO2.

### Immunofluorescence

Following sacrifice, bladders were removed and fixed in methacarn (60% methanol, 30% chloroform, and 10% acetic acid) for 20mins, before embedding within paraffin. The sections were deparaffinized by soaking in 3 separate solutions of 100% Histoclear for 5 minutes each. Sections were rehydrated in decreasing concentrations of ethanol (100%, 90%, 70%, 50%) for 5 minutes per concentration and soaked in 1X PBS for 5 minutes. Sections were blocked in 1% bovine serum albumin for 1 hour at room temperature, followed by antibody staining with antibodies against: Uroplakin III (Fitzgerald, NA), BiP (Cell Signaling Technology, C50B12), p27KIP1 (Abcam, ab190851), E-Cadherin (BD Biosciences 610181) and Ki67 (Abcam, ab833) in blocking buffer containing 0.1% Tween-20 overnight at 4°C. Sections were rinsed in 1X PBS (3 times, 5 minutes each), and incubated in appropriate fluorescently labeled secondary antibodies for 1 hour at room temperature and rinsed in 1X PBS (3 times, 5 minutes each). Sections were applied with Prolong Gold Antifade reagent with DAPI (P36935, Thermo Fisher Scientific, USA) and covered with cover glass before proceeding with imaging. Images were taken on either a Zeiss Apotome microscope at 40x magnification or ECLIPSE Ni Epi-fluorescence Upright Microscope (Nikon, USA).

### H&E staining

Bladder sections from WT and *Ifrd1^−/−^*mice were deparaffinized by soaking in 3 separate solutions of 100% Histoclear for 5 minutes each. Sections were rehydrated in decreasing concentrations of ethanol (100%, 90%, 70%, 50%) for 5 minutes per concentration. Staining of the nuclei was performed by soaking the sections in Hematoxylin solution for 5 minutes followed by rinsing in running tap water to remove excess Hematoxylin solution. Sections were dipped quickly into an acid alcohol solution, soaked for 5 minutes in sodium bicarbonate solution, followed by rinsing in distilled water. Staining of the cytosol was done immediately by dipping the slides 3 times in Eosin solution, soaking in increasing concentrations of ethanol (50%, 70%, 90%, 100%) for 3 minutes each, followed by a final dip in Histoclear. Sections were applied with xylene-based mounting medium, covered with cover glass, and edges were sealed with clear nail polish. Image capture was performed with a Panoramic Midi microscope (3DHISTECH Ltd, Hungary).

### Transmission electron microscopy (TEM) and quantification of cell structures and organelles

Whole bladders of WT and *Ifrd1^−/−^* mice were fixed using 2% glutaraldehyde and 3% paraformaldehyde in 0.1 M sodium cacodylate. Samples were then washed a total of three times in sodium cacodylate buffer and post-fixed in 1% osmium tetroxide for one hour, and then stained in 1% uranyl acetate for an hour, before being rinsed, dehydrated, and subjected to critical point drying. Bladder samples were then gold-coated and viewed using JEOL 1200 EX II Transmission Electron Microscope (JEOL, USA). For quantification of MVB, lysosomes, and mitochondria, images of superficial cells were taken at 2500x. The number of MVBs, cargo-containing MVBs, mitochondria, and lysosomes were quantified, and the amounts were reported as per 100µm^2^ surface area examined. (n= 50-150 TEM sections from four individual mice).

### TUNEL and Puromycylation assays

After sacrifice, mouse bladders were flash frozen using OCT. Seven-micron sections were then stained with TUNEL according to manufactures protocol (Roche 11684795910, In Situ Cell Death Detection Kit). Images were taken on a Zeiss Apotome microscope (n=4 mice). For puromycylation, the sections were fixed in 10% formalin for 15 min at RT, rinsed in 1X PBS (3 times, 5 minutes each) and then blocked in blocked in 1% bovine serum albumin containing 0.1% Triton X-100. Sections were then incubated overnight with antibody against puromycin (MABE343, EMD Millipore) at 4 °C. Sections were rinsed in 1X PBS (3 times, 5 minutes each), and incubated in fluorescently labeled secondary antibody for 1 hour at room temperature and rinsed in 1X PBS (3 times, 5 minutes each). Sections were applied with Prolong Gold Antifade reagent with DAPI (P36935, Thermo Fisher Scientific, USA) and covered with cover glass before proceeding with imaging. Images were taken on ECLIPSE Ni Epi-fluorescence Upright Microscope (Nikon, USA). For quantification of puromycin signal, mean intensity of 3-4 fields from 3-4 different 40X images were used (n=3 mice).

### Urine cytology and Dihydroethidium ROS Assay

WT and *Ifrd1^−/−^* mouse urine samples were collected for sediment analysis (10 μl urine plus 40 μl 1X PBS) and subjected to cytospin3. Sediments on the microscope slides were fixed in acetic acid/alcohol for 15 minutes and subjected to Epredia™ Papanicolaou EA Staining (22-050-211, Thermo Fisher Scientific, USA) following the manufacturer’s protocol. Bright field images from a whole slide scanner were used to identify and count the sloughed urothelial cells.

To assess for the production of ROS by the sloughed superficial cells, 50µl of urine was centrifuged to collect cells that were then incubated with 10μM dihydroethidium (ThermoFIsher Scientific, D23107) for 30 min at 37°C, rinsed in PBS, and then subjected to cytospin3. Subsequently they were mounted with Prolong Diamond Antifade Mountant with DAPI (ThermoFisher Scientific, P36962) and promptly imaged using Zeiss Axio Imager M2 Plus Wide Field Fluorescence Microscope.

### Co-Immunoprecipitation and Proteomic Analysis

Dynabeads Protein A for Immunoprecipitation (ThermoFisher, 10001D) were resuspended in PBS-Tween (0.2%)+ Protease Inhibitor, before 10min incubation with anti-IFRD1 (Abcam, ab229720) and Rabbit IgG (Cell Signaling, 2729S). Cultured 5637 human bladder cancer cells were lysed in Pierce IP Lysis Buffer (ThermoFisher, 87787), and incubated for 30mins at room temperature. Following centrifugation of the lysate, the supernatant was transferred to the bead now attached to the antibodies of interest. Beads were incubated overnight at 4 °C, and then gently washed to remove unbound proteins. Samples were then submitted to the Proteomics Core Laboratory at Washington University in St Louis for MS/MS analysis.

For database searching, the tandem mass spectra were extracted by the search engine Mascot (version 2.5.1). Mascot searched the UNI-HUMAN-REF-20190731 database containing 20667 entries. Mascot was searched with a fragment ion mass tolerance of 0.05.0 Da and a parent ion tolerance of 25 parts per million. In Mascot, Carbamidomethyl of cysteine was specified as a fixed modification; and the following were specified as variable modifications: Gln->pyro-Glu of the n terminus, deamidated of asparagine and glutamine, oxidation of methionine and acetyl of the n terminus.

Criteria for protein identification was done using Scaffold_4.11.0 Proteome Software, which validated MS/MS based peptide and protein identifications. Identification of peptides were accepted if established at >91.0% probability to achieve an FDR >1.0%, using the Scaffold Local FDR algorithm. Proteins were identified and accepted if established at >66.0% probability, with a minimum of 1 identified peptide. Proteins probabilities were assigned using the Protein Prophet algorithm (Adapted from Scaffold_4.11.0).

### RNA-Sequencing

Preparation of bladders for RNA isolation and sequencing were done as described in Ligon et al (2020). In short, bladders were snap frozen and homogenized for RNA isolation using RNeasy Mini Kit (Qiagen, 74101). Preparation of libraries was done with Ribo-Zero rRNA depletion kit (Illumina) and then sequenced on HiSeq3000 (Illumina). Reads were then aligned to the Ensemble top-level assembly with STAR version 2.0.4b. Gene counts were produced from the uniquely aligned unambiguous reads by Subread:feature Count version 1.4.5, and transcript counts were derived by Sailfish version 0.6.3. Using RSeQC version 2.3, sequencing performance was determined for the total number of aligned reads, total number of uniquely aligned reads, genes and transcripts detected, ribosomal fraction, known junction saturation and read distribution over known gene models. All gene counts were then imported into R/Bioconductor package EdgeR and TMM normalization size factors were calculated to adjust samples for differences in library size. Some features, such as ribosomal and any feature not expressed in at least three samples, were were excluded from further analysis and TMM size factors were recalculated to create effective TMM size factors. These factors and matrix counts were then imported into R/Bioconductor package Limma, and the voomWithQualityWeights function was used to calculate the observed mean-variance relationship of every gene/transcript and sample. Generalized linear models were then used to test for gene/transcript level differential expression, which were filtered for False Discovery Rate (FDR)-adjusted p-values less than or equal to 0.05.

### Quantitative RT-PCR (qRT-PCR)

Bladders of young WT and *Ifrd1^−/−^* mice were collected and homogenized in TRIzol™ Reagent (15596026, Thermo Fisher Scientific, USA) to extract total RNA followed by DNase 1 treatment (18068-015, Thermo Fisher Scientific, USA) following manufacturer’s protocol. One microgram of total RNA was utilized to perform cDNA synthesis using SuperScript™ II Reverse Transcriptase (18064-014, Thermo Fisher Scientific, USA) following manufacturer’s protocol. All cDNAs were diluted to 1:8 with RNase-free water prior thereafter. Primer designs (see Key Resources Table) and qRT-PCR setup was performed using SsoAdvanced Universal SYBR® Green Supermix (1725274, Bio-Rad, USA) following manufacturer’s protocol in 10 μl reactions (5 μl Supermix; 1 μl each of Forward and Reverse Primers; 2 μl diluted cDNA; 1 μl RNase-free water), each reaction was done in triplicate. 18s rRNA was used as a housekeeping gene. The qRT-PCR reaction was run in QuantStudio™3 Real-Time PCR System (Applied Biosystems™, USA) using the following settings: 98 °C, 3 minutes (initial activation); 98 °C, 30 seconds (Denaturation); 58 °C, 30 seconds (Annealing/Extension); 40 cycles; and instrument default setting for melt-curve analysis. Raw quantitation values were used for calculating fold change based on the WT bladder. See TABLE 1 for the sequences of primers used.

**TABLE 1:**
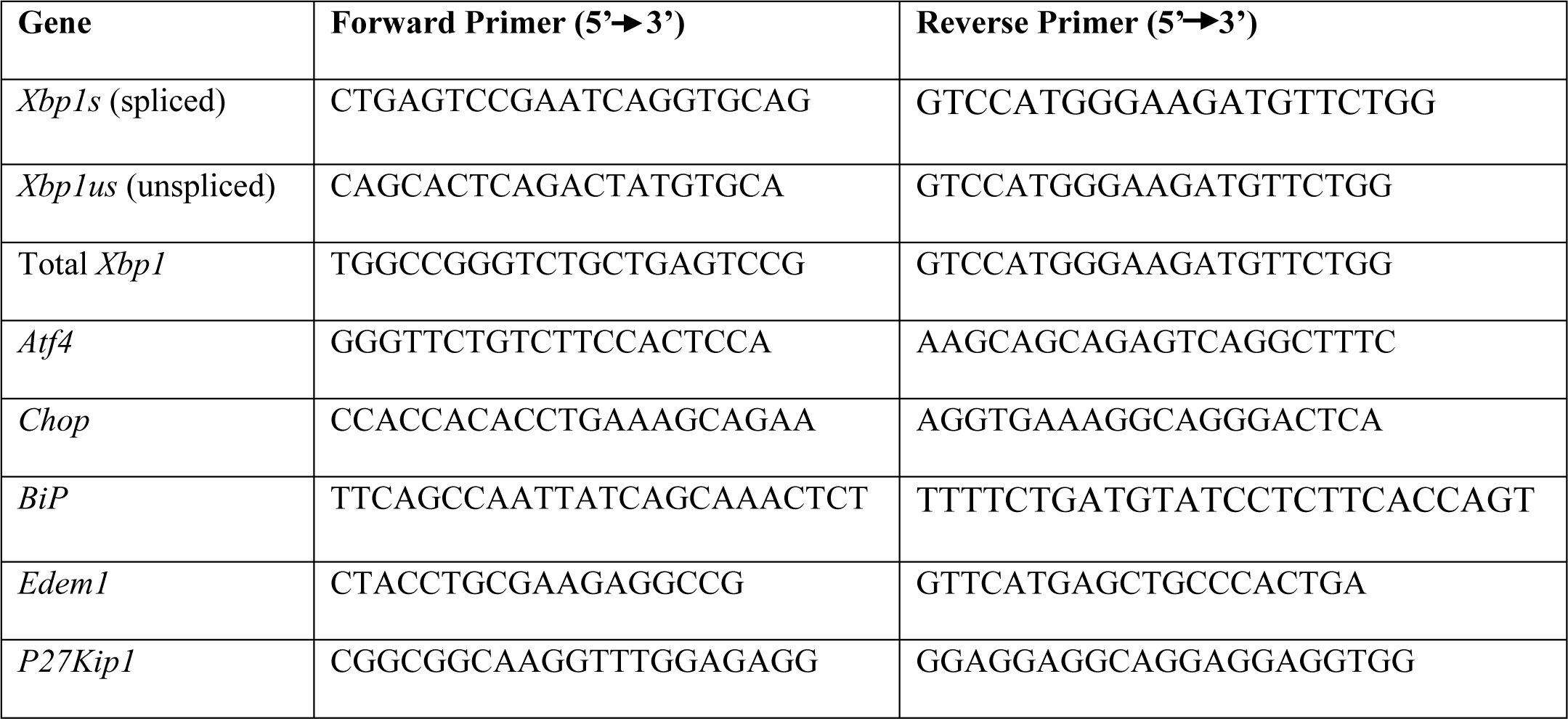
Primer sequences used in RT-qPCR analysis.

### Western Blotting

WT and *Ifrd1^−/−^* bladders of young mice were collected and homogenized in RIPA lysis buffer to extract total proteins. Total protein was quantitated using a BCA assay (Pierce™ BCA Protein Assay Kit, 23225, Thermo Fisher Scientific, USA). 10-30 µg of protein was loaded onto precast gel (NP0321, NuPAGE™ 4- 12%, Bis-Tris, Mini Protein Gels, ThermoFisher Scientific, USA) and resolved at 135 volts for 90 minutes. Proteins bands were immobilized through transfer to PVDF membrane (IPFL00010, Immobilon Transfer Membrane, Millipore, Ireland) for 90 min at 30 volts in ice. The membrane was treated with blocking buffer (927-60001, Intercept® (TBS) Blocking Buffer, LI-COR, USA) for 1 hour at room temperature with gentle agitation. Membranes were incubated with primary antibody [IFRD1 (abcam, ab229720), BiP (CST, 3177S), ATF6 (Thermo Fisher, PA5-20216), XBP1 (CST, 40435S), Edem1 (Proteintech, 26226-1-AP), PERK (CST, 5683S), p-PERK (Thermo Fisher, PA5-102853), eIF2α (CST, 9722S), p-eIF2α (CST, 3398S), ATF4 (Abcam, AB1371), CHOP (CST, 2895S)] solution (1:1000) in blocking buffer plus 0.1% Tween-20, overnight at 4°C with agitation. Beta-actin (CST, 3700S) was used as a loading control. Membranes were washed in 1X TBS with 0.1% Tween-20, 5 times, for 5 minutes each at room temperature with agitation followed by treatment with appropriate secondary antibody solution in blocking buffer plus 0.1% Tween-20 for 1 hour at room temperature with agitation. Membranes were washed in 1X TBS with 0.1% Tween-20, 5 times, for 5 minutes each at room temperature with agitation followed by washing in 1x TBS 2 times, for 5 minutes each. Imaging was performed using the ChemiDoc^TM^ MP imaging system (Bio-Rad, USA). Densitometric analysis was done using Bio-Rad Image Lab software (6.1).

### Statistics

All measured values were plotted using GraphPad Prism versions 9.0.1 to 10.0.3 (GraphPad Software, La Jolla, CA, USA). Data were expressed as mean ± Standard Deviation (SD) (indicated in each figure).

## Notes

### Competing Interest Statement

The authors have declared no competing interest.

## REFERENCES

1. Rajasekaran, M., Stein, P. & Parsons, C. L. Toxic factors in human urine that injure urothelium. Int J Urol 13, 409–414 (2006).

2. Hicks, R. M., Ketterer, B. & Warren, R. C. The ultrastructure and chemistry of the luminal plasma membrane of the mammalian urinary bladder: a structure with low permeability to water and ions. https://royalsocietypublishing.org/doi/epdf/10.1098/rstb.1974.0013 (1974) doi:10.1098/rstb.1974.0013.

3. Jost, S. P. Cell cycle of normal bladder urothelium in developing and adult mice. Virchows Archiv B Cell Pathol 57, 27–36 (1989).

4. Jost, S. P. & Potten, C. S. Urothelial Proliferation In Growing Mice. Cell Proliferation 19, 155–160 (1986).

5. Colopy, S. A., Bjorling, D. E., Mulligan, W. A. & Bushman, W. A population of progenitor cells in the basal and intermediate layers of the murine bladder urothelium contributes to urothelial development and regeneration. Developmental Dynamics 243, 988–998 (2014).

6. Gandhi, D. et al. Retinoid Signaling in Progenitors Controls Specification and Regeneration of the Urothelium. Developmental Cell 26, 469–482 (2013).

7. Mulvey, M. A. et al. Induction and evasion of host defenses by type 1-piliated uropathogenic Escherichia coli. Science 282, 1494–1497 (1998).

8. Mysorekar, I. U., Mulvey, M. A., Hultgren, S. J. & Gordon, J. I. Molecular regulation of urothelial renewal and host defenses during infection with uropathogenic Escherichia coli. J Biol Chem 277, 7412–7419 (2002).

9. Mysorekar, I. U., Isaacson-Schmid, M., Walker, J. N., Mills, J. C. & Hultgren, S. J. Bone morphogenetic protein 4 signaling regulates epithelial renewal in the urinary tract in response to uropathogenic infection. Cell Host Microbe 5, 463–475 (2009).

10. Romih, R., Koprivec, D., Martincic, D. S. & Jezernik, K. Restoration of the rat urothelium after cyclophosphaheximide after treatment. Cell Biology International 25, 531–537 (2001).

11. Hyuga, T. et al. Wound healing responses of urinary extravasation after urethral injury. Sci Rep 13, 10628 (2023).

12. Jafari, N. V. & Rohn, J. L. The urothelium: a multi-faceted barrier against a harsh environment. Mucosal Immunol 15, 1127–1142 (2022).

13. Kreft, M. E., Sterle, M., Veranič, P. & Jezernik, K. Urothelial injuries and the early wound healing response: tight junctions and urothelial cytodifferentiation. Histochem Cell Biol 123, 529–539 (2005).

14. Wang, C., Ross, W. T. & Mysorekar, I. U. Urothelial Generation and Regeneration in Development, Injury, and Cancer. Dev Dyn 246, 336–343 (2017).

15. Balestreire, E. M. & Apodaca, G. Apical epidermal growth factor receptor signaling: regulation of stretch-dependent exocytosis in bladder umbrella cells. Mol Biol Cell 18, 1312–1323 (2007).

16. Chen, Y. et al. Rab27b is associated with fusiform vesicles and may be involved in targeting uroplakins to urothelial apical membranes. Proc Natl Acad Sci U S A 100, 14012–14017 (2003).

17. Lewis, S. A. & de Moura, J. L. Incorporation of cytoplasmic vesicles into apical membrane of mammalian urinary bladder epithelium. Nature 297, 685–688 (1982).

18. Minsky, B. & Chlapowski, F. Morphometric analysis of the translocation of lumenal membrane between cytoplasm and cell surface of transitional epithelial cells during the expansion-contraction cycles of mammalian urinary bladder. Journal of Cell Biology 77, 685–697 (1978).

19. Truschel, S. T. et al. Stretch-regulated exocytosis/endocytosis in bladder umbrella cells. Mol Biol Cell 13, 830–846 (2002).

20. Amano, O., Kataoka, S. & Yamamoto, T. Y. Turnover of asymmetric unit membranes in the transitional epithelial superficial cells of the rat urinary bladder. Anat Rec 229, 9–15 (1991).

21. Bäck, N., Rajagopal, C., Mains, R. E. & Eipper, B. A. Secretory Granule Membrane Protein Recycles through Multivesicular Bodies. Traffic 11, 972–986 (2010).

22. Khandelwal, P., Ruiz, W. G. & Apodaca, G. Compensatory endocytosis in bladder umbrella cells occurs through an integrin-regulated and RhoA- and dynamin-dependent pathway. EMBO J 29, 1961–1975 (2010).

23. Zhou, G. et al. MAL facilitates the incorporation of exocytic uroplakin-delivering vesicles into the apical membrane of urothelial umbrella cells. Mol Biol Cell 23, 1354–1366 (2012).

24. Iezaki, T. et al. Transcriptional Modulator Ifrd1 Regulates Osteoclast Differentiation through Enhancing the NF-κB/NFATc1 Pathway. Mol Cell Biol 36, 2451–2463 (2016).

25. Park, G. et al. The transcriptional modulator Ifrd1 controls PGC −1α expression under short-term adrenergic stimulation in brown adipocytes. The FEBS Journal 284, 784–795 (2017).

26. Tummers, B. et al. The interferon-related developmental regulator 1 is used by human papillomavirus to suppress NFκB activation. Nat Commun 6, 6537 (2015).

27. Miao, Z.-F. et al. A Dedicated Evolutionarily Conserved Molecular Network Licenses Differentiated Cells to Return to the Cell Cycle. Developmental Cell 55, 178–194.e7 (2020).

28. Willet, S. G. et al. Regenerative proliferation of differentiated cells by mTORC1-dependent paligenosis. EMBO J 37, e98311 (2018).

29. Brown, J. W., Cho, C. J. & Mills, J. C. Paligenosis: Cellular Remodeling During Tissue Repair. Annu. Rev. Physiol. 84, 461–483 (2022).

30. Andreev, D. E. et al. Translation of 5’ leaders is pervasive in genes resistant to eIF2 repression. Elife 4, e03971 (2015).

31. Zhao, C., Datta, S., Mandal, P., Xu, S. & Hamilton, T. Stress-sensitive regulation of IFRD1 mRNA decay is mediated by an upstream open reading frame. J Biol Chem 285, 8552–8562 (2010).

32. Schmidt, E. K., Clavarino, G., Ceppi, M. & Pierre, P. SUnSET, a nonradioactive method to monitor protein synthesis. Nat Methods 6, 275–277 (2009).

33. Read, A. & Schröder, M. The Unfolded Protein Response: An Overview. Biology 10, 384 (2021).

34. Walter, P. & Ron, D. The unfolded protein response: from stress pathway to homeostatic regulation. Science 334, 1081–1086 (2011).

35. Haze, K., Yoshida, H., Yanagi, H., Yura, T. & Mori, K. Mammalian transcription factor ATF6 is synthesized as a transmembrane protein and activated by proteolysis in response to endoplasmic reticulum stress. Mol Biol Cell 10, 3787–3799 (1999).

36. Park, S.-M., Kang, T.-I. & So, J.-S. Roles of XBP1s in Transcriptional Regulation of Target Genes. Biomedicines 9, 791 (2021).

37. Gordiyenko, Y., Llácer, J. L. & Ramakrishnan, V. Structural basis for the inhibition of translation through eIF2α phosphorylation. Nat Commun 10, 2640 (2019).

38. Kimball, S. R., Fabian, J. R., Pavitt, G. D., Hinnebusch, A. G. & Jefferson, L. S. Regulation of guanine nucleotide exchange through phosphorylation of eukaryotic initiation factor eIF2alpha. Role of the alpha- and delta-subunits of eiF2b. J Biol Chem 273, 12841–12845 (1998).

39. Nishitoh, H. CHOP is a multifunctional transcription factor in the ER stress response. J Biochem 151, 217–219 (2012).

40. Jost, S. P. Renewal of normal urothelium in adult mice. Virchows Arch B Cell Pathol Incl Mol Pathol 51, 65–70 (1986).

41. Papafotiou, G. et al. KRT14 marks a subpopulation of bladder basal cells with pivotal role in regeneration and tumorigenesis. Nat Commun 7, 11914 (2016).

42. Arenander, A. T. et al. TIS gene expression in cultured rat astrocytes: induction by mitogens and stellation agents. J Neurosci Res 23, 247–256 (1989).

43. Guardavaccaro, D., Ciotti, M. T., Schäfer, B. W., Montagnoli, A. & Tirone, F. Inhibition of differentiation in myoblasts deprived of the interferon-related protein PC4. Cell Growth Differ 6, 159–169 (1995).

44. Vadivelu, S. K. et al. Muscle Regeneration and Myogenic Differentiation Defects in Mice Lacking TIS7. Mol Cell Biol 24, 3514–3525 (2004).

45. Wang, Y. et al. Targeted intestinal overexpression of the immediate early gene tis7 in transgenic mice increases triglyceride absorption and adiposity. J Biol Chem 280, 34764–34775 (2005).

46. Nelson, D. P. et al. Myocardial immediate early gene activation after cardiopulmonary bypass with cardiac ischemia-reperfusion. The Annals of Thoracic Surgery 73, 156–162 (2002).

47. Roth, A., Gill, R. & Certa, U. Temporal and spatial gene expression patterns after experimental stroke in a rat model and characterization of PC4, a potential regulator of transcription. Mol Cell Neurosci 22, 353–364 (2003).

48. Rubin, D. C., Swietlicki, E. A., Wang, J. L. & Levin, M. S. Regulation of PC4/TIS7 expression in adapting remnant intestine after resection. American Journal of Physiology-Gastrointestinal and Liver Physiology 275, G506–G513 (1998).

49. Radyk, M. D. et al. ATF3 induces RAB7 to govern autodegradation in paligenosis, a conserved cell plasticity program. EMBO Rep 22, e51806 (2021).

50. Schröder, M. & Kaufman, R. J. The mammalian unfolded protein response. Annu Rev Biochem 74, 739–789 (2005).

51. Zhang, K. & Kaufman, R. J. From endoplasmic-reticulum stress to the inflammatory response. Nature 454, 455–462 (2008).

52. Ligon, M. M., Joshi, C. S., Fashemi, B. E., Salazar, A. M. & Mysorekar, I. U. Effects of aging on urinary tract epithelial homeostasis and immunity. Developmental Biology 493, 29–39 (2023).

53. Mulvey, M. A., Schilling, J. D., Martinez, J. J. & Hultgren, S. J. Bad bugs and beleaguered bladders: interplay between uropathogenic Escherichia coli and innate host defenses. Proc Natl Acad Sci U S A 97, 8829–8835 (2000).

54. Joshi, C. S. et al. D-Mannose reduces cellular senescence and NLRP3/GasderminD/IL-1β-driven pyroptotic uroepithelial cell shedding in the murine bladder. Dev Cell S1534-5807(23)00615–9 (2023) doi:10.1016/j.devcel.2023.11.017.

55. Micheli, L. et al. PC4 coactivates MyoD by relieving the histone deacetylase 4-mediated inhibition of myocyte enhancer factor 2C. Mol Cell Biol 25, 2242–2259 (2005).

56. Vietor, I. et al. TIS7 interacts with the mammalian SIN3 histone deacetylase complex in epithelial cells. EMBO J 21, 4621–4631 (2002).

57. Vietor, I., Kurzbauer, R., Brosch, G. & Huber, L. A. TIS7 regulation of the beta-catenin/Tcf-4 target gene osteopontin (OPN) is histone deacetylase-dependent. J Biol Chem 280, 39795–39801 (2005).

58. Balci, M. G. & Tayfur, M. Loss of E-cadherin expression in recurrent non-invasive urothelial carcinoma of the bladder. Int J Clin Exp Pathol 11, 4163–4168 (2018).

59. Zhao, Y., Xia, Z., Lin, T. & Yin, Y. Significance of hub genes and immune cell infiltration identified by bioinformatics analysis in pelvic organ prolapse. PeerJ 8, e9773 (2020).

60. Zhou, Y., Liu, X., Li, W., Sun, X. & Xie, Z. Endoplasmic reticulum stress contributes to the pathogenesis of stress urinary incontinence in postmenopausal women. J Int Med Res 46, 5269– 5277 (2018).

61. Alperin, M. et al. Foundational Science and Mechanistic Insights for a Shared Disease Model: An Expert Consensus. Female Pelvic Med Reconstr Surg 28, 347–350 (2022).

